# The prokaryotic SPHINX 1.8 REP protein is tissue-specific and expressed in human germline cells

**DOI:** 10.1101/373803

**Authors:** Laura Manuelidis

**Affiliations:** Yale University Medical School, Section of Neuropathology, Surgery Department, 333 Cedar Street, New Haven, CT 06510

## Abstract

Small circular DNAs of 1.8 and 2.4kb were initially discovered in highly infectious CJD and scrapie particles from mammalian brain and cultured cells. Surprisingly, these protected cytoplasmic "SPHINX" DNAs contained replication (REP) initiation sequences resembling those of Acinetobacter phage viruses. An antibody was generated against a REP peptide encoded by the SPHINX 1.8 ORF that was not present in mammals. It bound to a 41kd "spx1" protein on Western blots. Cytologically, spx1 concentrated in spinal cord synapses and pancreatic islet, but not exocrine cells. We hypothesized that circular SPHINX DNAs are ancient symbiotic elements that can participate in functional differentiation and neurodegeneration. Cell and tissue specific patterns of spx1 expression shown below implicate somatic cell-to-cell communication and differentiation functions that would favor conservation of SPHINX 1.8 in evolution. Remarkably, primary human oocytes and spermatogonia, but not mature sperm, displayed intense cytoplasmic spx1 signals that underscore the maternal inheritance of SPHINX 1.8. These findings should encourage investigations of unexplored networks of incorporated environmental infectious agents that can be key actors in progressive neurodegeneration, immunity and cancer.

## INTRODUCTION

Transmissible encephalopathies (TSEs) such as infectious human Creutzfeldt-Jakob Disease (CJD), endemic sheep scrapie, and epidemic bovine encephalopathy (BSE), show many progressive features and convergent markers with Alzheimer’s Disease (AD) and other "non-infectious" neurodegenerative disorders. Such features include accumulation of host-encoded amyloids and various fibril aggregates, microglial activation, and molecular markers such as Clusterin (Apo J) and MIP1*α* (1-4). Therapeutic targeting of AD amyloid in humans has repeatedly failed, even with drugs that reduce amyloid plaques (5), because *β*-amyloid, as prion protein (PrP) amyloid, is not the initiating cause of disease, but rather, a late stage pathological product (1, 6-9). Large amounts of amyloid injected into the brain will collect as deposits that can be toxic to neurons in AD but are not transmissible in serial passages (10), and the popular belief that recombinant PrP (recPrP) amyloid is infectious is based on a single brain homogenate from very old transgenic mice that were inoculated with large amounts of "seeded" recPrP amyloid fibrils. These "seeded" mice transmitted only laboratory RML scrapie on serial passages (11). Independent investigators have not been able to generate infectivity from recPrP itself despite many robust attempts, e.g., (7, 8, 12). Nevertheless, amyloid may have other important functions. In fact, amyloid has been shown to function as an innate immune trap that can clear TSE infectious agents (1, 13), and it may also ensnare specific cells and clearance molecules. Many viruses also possess amyloid domains with critical functions such as viral transcription and receptor-mediated attachment to the host cell (14); the latter interaction fits neatly with the concept that host PrP is an essential receptor for an environmental TSE virus (1). While proteomics has highlighted common cellular proteins in neurodegeneration e.g., (6, 15), it may overlook past or covert causal infectious agents, especially those that collaborate with human endogenous viruses.

The prion seeding model for amyloids and other fibrillar accumulations in late onset human neurodegenerative diseases has largely discounted causal environmental pathogens. Our laboratory has been particularly intrigued by the extreme latency of TSE infections and their viral strain-specific phenotypes (1, 16). Highly infectious 20-25nm dense particles isolated from brain and cultured cells correspond to those observed ultrastructurally within TSE infected, but not control cells (6, 17, 18), and TSE agents may link to a larger group of covert, incorporated environmental agents involved in some forms of AD, Parkinson’s disease (PD) and other progressive neurodegenerations. Established viruses are known to elicit common neuropathological stigmata, and the 1918 influenza epidemic caused late-onset PD many years after the virus was eliminated from the host. Moreover, post-encephalitic PD, as AD, is manifest by classical neurofibrillary tangles, and these stigmata are also induced by subacute myxo-paramyxoviral infections. Other commensal viruses, such as Herpes, have also been implicated in AD (19), and only now are receiving renewed attention (20). A role for new or unrecognized viral species also deserves consideration.

The biologic and physical viral properties of infectious TSE particles (18) led to the discovery of new unsuspected SPHINX DNAs. Small circular DNAs of 1.5-4kb, such as those in 17-22nm circoviruses, fit the experimental size of TSE infectious particles. Given the extreme stability of individual TSE agent strains (1), a DNA genome would be more likely than a rapidly mutating RNA. I therefore focused on unknown classes of circular DNAs in cytoplasmic infectious TSE particles using Phi29 polymerase for rolling circle DNA amplification (21). Such circular DNAs might be the infectious TSE agent itself, might act as satellite-associated collaborators for infection, or might function as key players in neurodegenerative processes. Two circular SPHINX DNAs were first isolated from scrapie and CJD infected hamster and mouse brain particles at much higher levels than in normal controls, and they were also readily detected in high titer nuclease protected particles from infected cell cultures (21). The 1.76kb and 2.36kb SPHINX DNAs were isolated, sequenced and deposited in the database in 2010 along with proof of their closed circular structure, upstream iteron sequences and ORFs with loose homology to phage replication (REP) initiator sequences. Although nucleases that destroy TSE particle infectivity also destroy these SPHINX DNAs (9), subsequent PCR studies demonstrated both circular DNAs in unfractionated cytoplasm of normal brain and cell cultures. Thus factors other than the presence of these two DNAs would be needed to produce phenotypic TSE infection. On the other hand, the same 1.76 SPHINX DNA (SPX1.8) was subsequently isolated from two European multiple sclerosis brains (22), strengthening a potential role for this circular DNA in neurodegeneration.

To the best of my knowledge, SPHINX DNAs are the only phage REPs reported to reside in the cytoplasm of mammalian cells. REP assemblies fall within the general class of circular single stranded "CISS" viruses that have been found within plant and invertebrate cells (23). Endogenous retroviruses (LINES), first isolated and sequenced in 1982 (24) derive from retroviruses, constitute as much as ~10% of nuclear DNA, and exemplify viral RNA sequences inserted as DNA copies into mammalian genomes early in evolution. These retroviral elements organize in large tissue and species specific chromosomal domains (25, 26) and can also be advantageous for selective survival. Xenotropic retroviruses of mice also advantageously cause disease only in competing foreign mice, a function that has relevance for the cross-species spread of HIV. Although contemporary DNA viruses that are acquired during one’s lifetime can also insert into nuclear DNA, unlike retroviruses they have not integrated into the germline. There has also been a flurry of recent interest in extrachromosomal circular DNAs in genome plasticity and disease (27-29), but these circular DNAs are comparatively huge (up to 16kb), and unlike SPHINX elements they do not contain prokaryotic sequences. Prior to the discovery of SPHINX DNAs only one known prokaryotic genome has been maintained in the cytoplasm of mammals. This is mitochondrial DNA (mtDNA), a protected circular DNA of 16kb that is maternally inherited through generations. It was originally derived by endosymbiosis from a bacterial genome, as first brilliantly deduced by Lynn Margulis (30). Its genome provides many metabolic functions that ensure its continued propagation in mammals.

To further explore the possible function(s) of SPX1.8 in brain and other tissues, antibodies were raised in rabbits against two peptides with no homology to any mammalian sequence in the database (31). Both rabbit antibodies bound to a 41kd non-glycosylated "spx1" protein band on Western blots, in reasonable accord with the predicted 328aa REP transcript located immediately downstream from its classical prokaryotic iteron (21). One rabbit antisera was affinity purified and its binding to spx1 was completely blocked by the prokaryotic antigenic REP peptide. This antibody was used to evaluate rodent brains, human glioblastoma cells, and pancreatic tissue (31). In all Western blot samples the 41kDa spx1 species was unambiguous. Normal rodent brains and those infected with CJD and scrapie, as well as PrP-null and 8x PrP producing mouse brains, all showed comparable levels of spx1 protein. Human glioblastoma samples also showed the same 41kd protein, indicating it is conserved in mammalian evolution. Selected in situ tissue studies further demonstrated a high concentration of the spx1 protein at synaptic boutons of the spinal cord. In addition, spx1 specifically decorated insulin secreting pancreatic islet cells but not exocrine cells. These cytologic studies again showed complete obliteration of signal by the cognate protein, and no spx1 signal was seen when the primary antibody was omitted. Together, this data strongly suggested a role for spx1 in differentiation, one that might ensure the propagation of its circular genome in a variety of mammals. The SPX1.8 DNA origin of the spx1 protein was further confirmed elsewhere in cells transfected with the full 1.8 REP aa sequence. Again, a prominent 41kDa Western blot band was highlighted using that antibody raised against the whole REP protein (32).

We here report extensive cytological analyses of various tissues using our spx1 antibody to interrogate various peripheral tissues from hamsters, mice, rats and humans. These studies demonstrate the differential expression, progressive development, and concentration of spx1 only in selected cell types. The patterns of staining implicate particular functions. Additionally, the strong germline expression of spx1 in primary oocytes and spermatogonia of humans underscores the incorporation of SPX1.8 into the germline of mammals in the distant past, a finding is not compatible with a recent environmental or artifactual contaminant. This germline residence of spx1, and its loss in mature sperm is also most consistent with a maternal inheritance of SPX1.8. A separate paper will detail the conserved synapse-specific patterns of spx1 across mammalian species. The role(s) of spx1 in differentiation, aging, and neoplastic transformation present intriguing new avenues for evolutionary, microbiological and functional inquiries.

## RESULTS

The following results are based on >5 samples from different mice, a minimum of 3 from rats, germinal human tissue (3 testis and 2 ovary), and a variety of human skin biopsy samples. Representative archival paraffin-embedded section were dewaxed and either not pretreated, or were exposed for 10 min to 2M GdnHCl to increase spx1 antigen unmasking prior to antibody detection as previously described in detail (31). Results were typically indistinguishable without and with GdnHCl as shown in Fig. 1A. Strong red signals are seen in both sets of the gastrointestinal (GI) tract. As in previous controls, the spx1 antigenic peptide completely blocked the signal and only the light blue hematoxylin counterstain is seen with negligible background. Pretreatment by citrate-heat unmasking (2) was also typically comparable as demonstrated below (see Fig. 4G & H).

**Fig 1:**
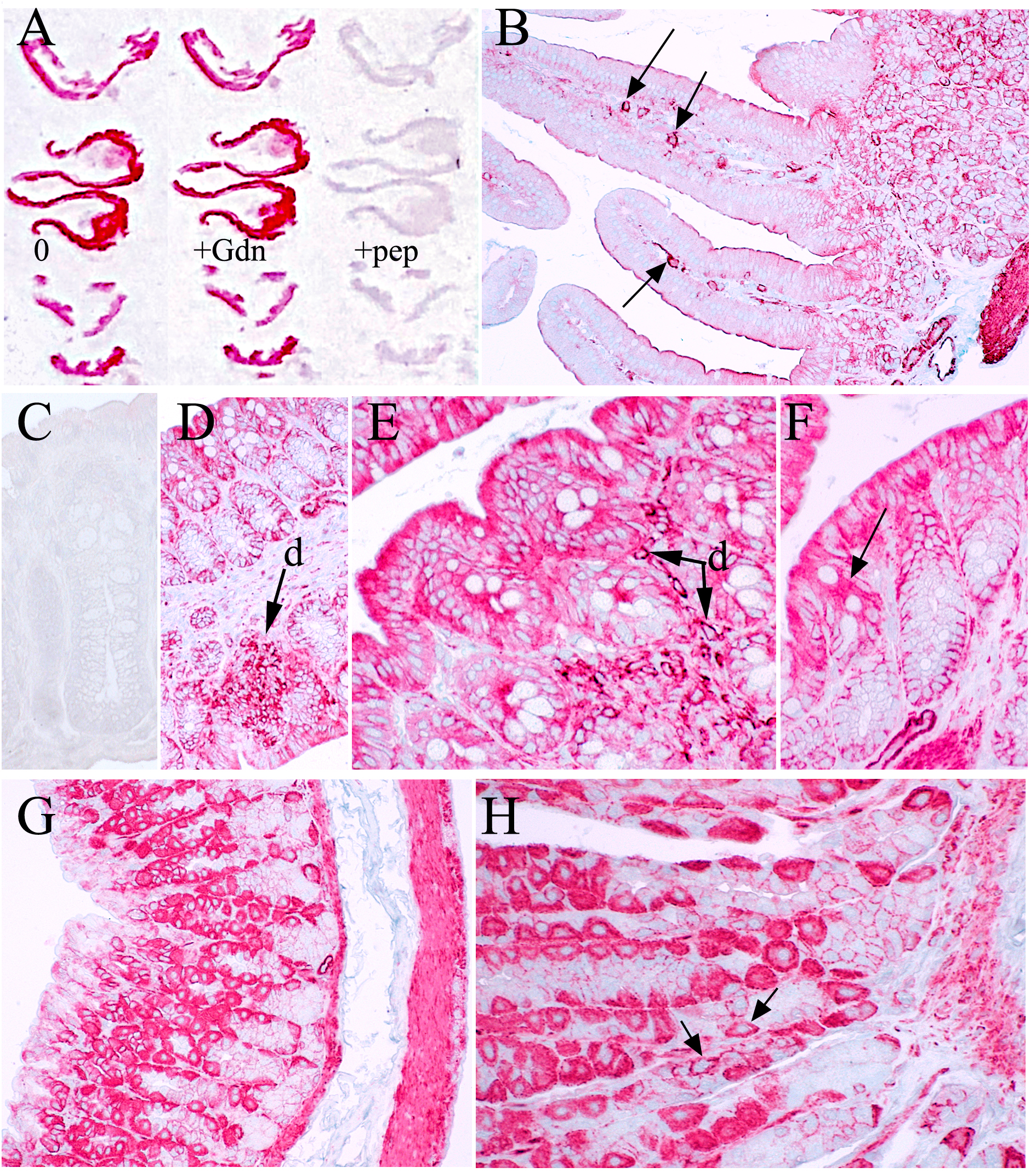
Affinity purified antibodies to spx1 detected in specific cells of mouse intestines. **A)** shows 3 parallel columns from 3 slides exposed to antibody after no pretreatment (0), after 2MGdnHCl, and after blocking with spx1 peptide (+pep). Equivalent strong red alkaline phosphatase is detected after 0 and 2M GdnHCl, whereas spx1 signals are completely blocked by peptide. **B)** Jejunum-Ileum mucosa shows minimal staining of cytoplasm with strong positive surface brush border. Lamina propria dendritic cells (arrows) are also strongly positive. **C-F)** show colon sections with peptide blocked control (**C**). In contrast, mucosa in a colon gland with lumen at top and bottom (**D)** also shows strong staining of cells with dendritic morphology in a large cluster at arrow. **E** show higher power of these cells. There is very strong cytoplasmic signal of goblet cells (arrow in **F**) and the surface epithelium shows more spx1 than more basilar cells, indicating progressive spx1 expression. **G &H)** Gastric mucosa without or with heat-citrate unmasking as described (2, 31) is the same. In both, chief and parietal cells and smooth muscle layers are strongly stained whereas superficial epithelium is minimally positive. In higher magnification **H**, minimally stained crypt cells show highly concentrated signal only at cell-to-cell contacts. Some small positive cells are probably neuroendocrine (arrows).

There was a marked difference in the intensity of spx1 protein in various cell types and regions of the GI tract. Fig. 1B shows only faint cytoplasmic staining of villous epithelial cells in the jejunum-ileum whereas dendritic cells (arrows) and the surface brush border are clearly positive. Intensely stained smooth muscle in this section, and in other peripheral tissue sections, provided an excellent internal positive control for negative cell types. Colon epithelium, in contrast to small intestine, displayed intense staining of the top layers (Fig. 1E), and spx1 was especially intense in well-developed goblet cells (F at arrow). A change in spx1 expression with cell-type differentiation is supported by a goblet cell cytoplasmic signal that becomes much more intense overall in superficial layer as compared to cells at the base. Dendritic-like cells (arrows at d in D & E) are again strongly positive, and blocking peptide abolished positive signals (compare C with equivalent section D). The increasing intensity of spx1 with differentiation from basal to more specialized goblet cells was further substantiated by sections of stomach (G & H). Extremely intense staining of chief and parietal cells contrasted with much weaker staining of surface neck cells and basal crypt cells. Moreover, basal cells displayed a different cell-to-cell plasma membrane pattern of staining rather than a diffuse cytoplasmic signal, consistent with different functional roles for spx1 in each of these cell types.

Sections of kidney (Fig. 2A) show a lack of detectable spx1 signal in the essentially passive filtration system of glomerular fenestrated capillaries and mesangial cells (at gl). Actively resorbing proximal convoluted tubules (pc) show a highly cell-specific pattern of intense brush border (surface cilia) staining in addition to some basal signal. In contrast, the proximal tubules (p) and distal tubules (d) show a more generalized intense cytoplasmic signal that is strongest at the base of the cells. In the kidney medulla (Fig. 2B), tubules also show strongly positive spx1 throughout the cytoplasm. Collecting duct cells (c) display an intermediate spx1 signal that is greater than negative glomerular cells and less intense than proximal tubule cells. In sum, spx1 shows a gradient of staining intensity that parallels positive ATP requiring active cells.

**Fig.2:**
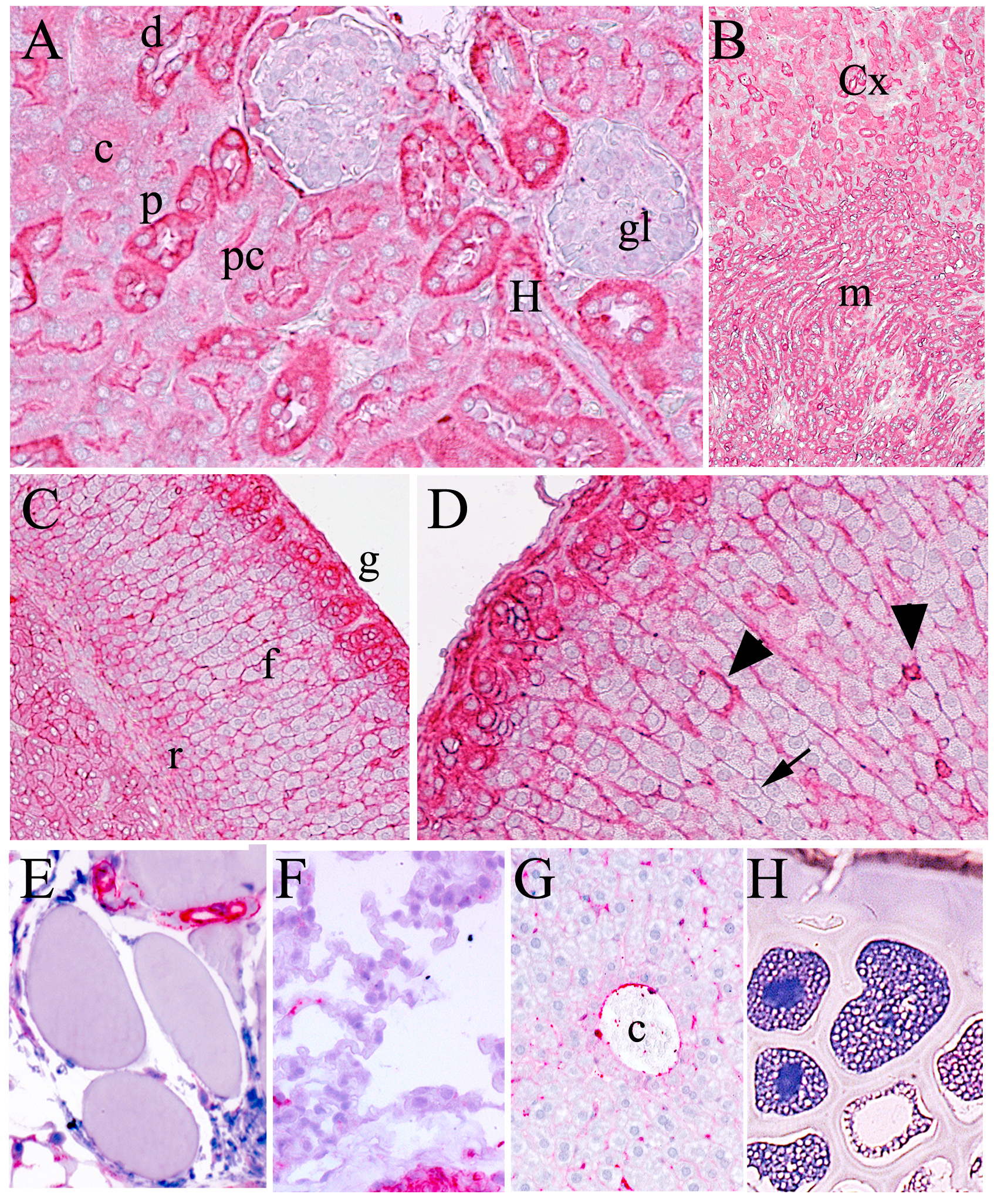
**A)** Mouse kidney cortex shows strong spx1 signal in selectively absorptive and secretory cells including proximal (p) and distal tubules (d) and Henle’s loop (H). Note intense staining of microvilli of brush border of proximal convoluted tubule (pc) with less intense cytoplasmic stain. In contrast to tubular epithelium, mesangial cells and fenestrated capillaries of glomeruli (gl) that basically act as passive filters are spx1 negative. A collecting duct cell (c) shows diffuse weak staining. **B)** shows low magnification of kidney cortex (Cx) and medulla (m) with intensely spx1 positive tubules. **C & D)** demonstrate major differences in spx1 protein in different adrenal cortex layers; staining is indistinguishable with 0 pretreatment versus GdnHCl pretreatment respectively. The outer glomerulosa layer (g) that manufactures aldosterone is strongly positive whereas the fasciculata that produces glucocorticoids shows negligible staining (arrow) where only thin fibers emanating from small mesenchymal cells are positive (arrowheads). Androgen producing cells of the reticularis (r) are moderately stained. Other cell types are essentially negative, including striated muscle **(E)**, lung **(F)**, hepatocytes **(G)** surrounding a central vein (c), and bone **(H)**.

The adrenal gland (Fig. 2C & D) also shows a remarkable differentiation of spx1 in functionally distinct cortical layers. The outer glomerulosa (g) layer produces the hormone aldosterone that induces resorption of sodium ions, and is intensely stained. In stark contrast, the glucocorticoid producing fasciculata is only marginally stained as shown in both GdnHCl pretreated and non-pretreated samples (2C & D respectively). Cords of fasciculata cells with minimal spx1 and are separated by thin positive strands of mesenchymal cells. The spx1 distinction between the two adrenal layers is notable because the positive glomerulosa cells with small mitochondria can be reversibly induced by ACTH to generate the enormous high ATP producing fasciculata mitochondria packed with elaborated cristae (33). Whereas spx1 increased with mitochondrial ATP activity in kidney, the adrenal showed the reverse pattern, i.e., fasciculata cells with high ATP activity mitochondria had low spx1. This indicates that other differentiated cellular properties control spx1 production.

Many tissues were essentially negative for spx1 including striated muscle (Fig. 2E), lung alveoli (F), hepatocytes (G), and bone (H) while smooth muscle and other cell types were strongly positive in these tissue sections. The spleen presented a more complex picture with positive and negative regional cell types, but precise spleen cell identities need more definitive combined marker studies for distinguishing related cell types. Nonetheless, there was a consistent overall pattern of intensely positive marginal zone cells (Fig. 3A at m). The splenic marginal zone is important in B-cell shuttling and follicular B-cell egress (34). In comparison, the red pulp has very sparse signal except for strongly positive megakaryocytes (arrows). Fig. 3C shows these large cells in another mouse spleen at higher power. The central lymphoid follicles (Fig. 3B) show positive dendritic-like cells. Fig. 3D shows spleen from an sCJD infected rat. The small lymphocytes show negligible spx1 whereas cells with an FDC morphology are positive (arrows) and FDC cells engage in TSE infections but are not required (35). The spx1 staining in spleen is comparable with GdnHCl unmasking (A & B) and without pretreatment (C & D). TSE infection and advanced age appeared to modify the number of selected spx1 positive selected cells such as FDC and this deserves more intense investigation with functional and sorting studies. Finally, skin showed strong spx1 signals at the border of hyperplastic squamous cells as shown in a human papilloma (verruca vulgaris) in Fig. 3E. At higher power there is bridge-like punctate desmosome pattern of labeling. Ascending squamous cells furthermore show more intense plasma membrane staining than basal cells, indicating accumulating spx1 with progressive differentiation in skin.

**Fig. 3:**
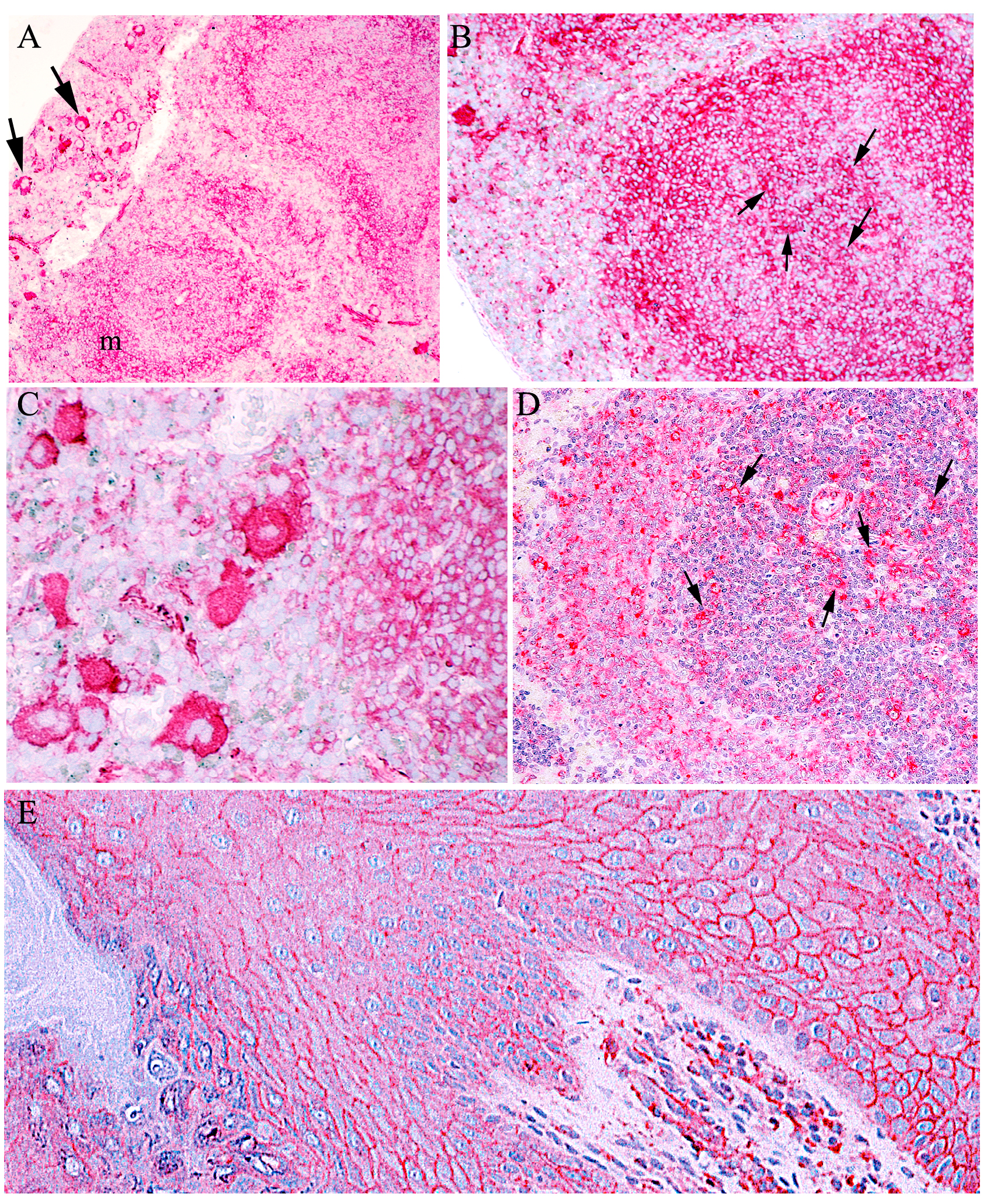
**A-C** Mouse spleens show a consistent strong staining pattern of the marginal zone that processes specialized B cells (34); the red pulp shows only a few positive. cell There is also an obvious and intense spx1 signal in large megakaryocytes in this non-hematoxylin counterstained GdnHCl pretreated section (arrows in A). Strong staining of non-lymphocyte cells in the lymphoid nodule (arrows in B) of a mouse, and C shows a higher power image of strongly spx1 positive megakaryocytes without GdnHCl pretreatment. **D)** shows no spx1 signal in lymphocytes, whereas an older sCJD infected rat shows strongly positive cells with an FDC morphology in the white pulp (arrows). **E)** Human papilloma (verruca vulgaris) section shows weakly positive cuboidal basal cells giving rise to stratified squamous cells. The prickle cell squamous layer show shows intensely spx1 positive plasma membranes in a dotted pattern consistent with concentration at desmosomal junctions. Stratum corneum is negative.

The testis showed the opposite developmental agenda with a progressive loss of spx1 from a strong signal in primary spermatogonia in seminiferous tubules that reside on the rim (Fig. 4A). Secondary spermatogonia are still positive but less intensely stained (arrows, 4B), whereas Sertoli cells (arrow st) and a Leydig cell (L) show weaker and negligible spx1 signals. Almost mature spermatocytes with some residual cytoplasm display only very weak signals indicating a progressive loss of spx1 with sperm maturation. In Fig. 4C, the arrow points to a cluster that includes labeled secondary spermatocytes that are larger (diploid) along with smaller (haploid) sperm, and both show detectable spx1. In contrast, sperm with minimal cytoplasm have negligible signal, and mature sperm that have lost their all mtDNA (at sp) have no spx1 signal. This picture is consistent with progressive loss and final elimination of SPX1.8 in mature sperm, and not merely a loss of its protein product.

**Fig.4:**
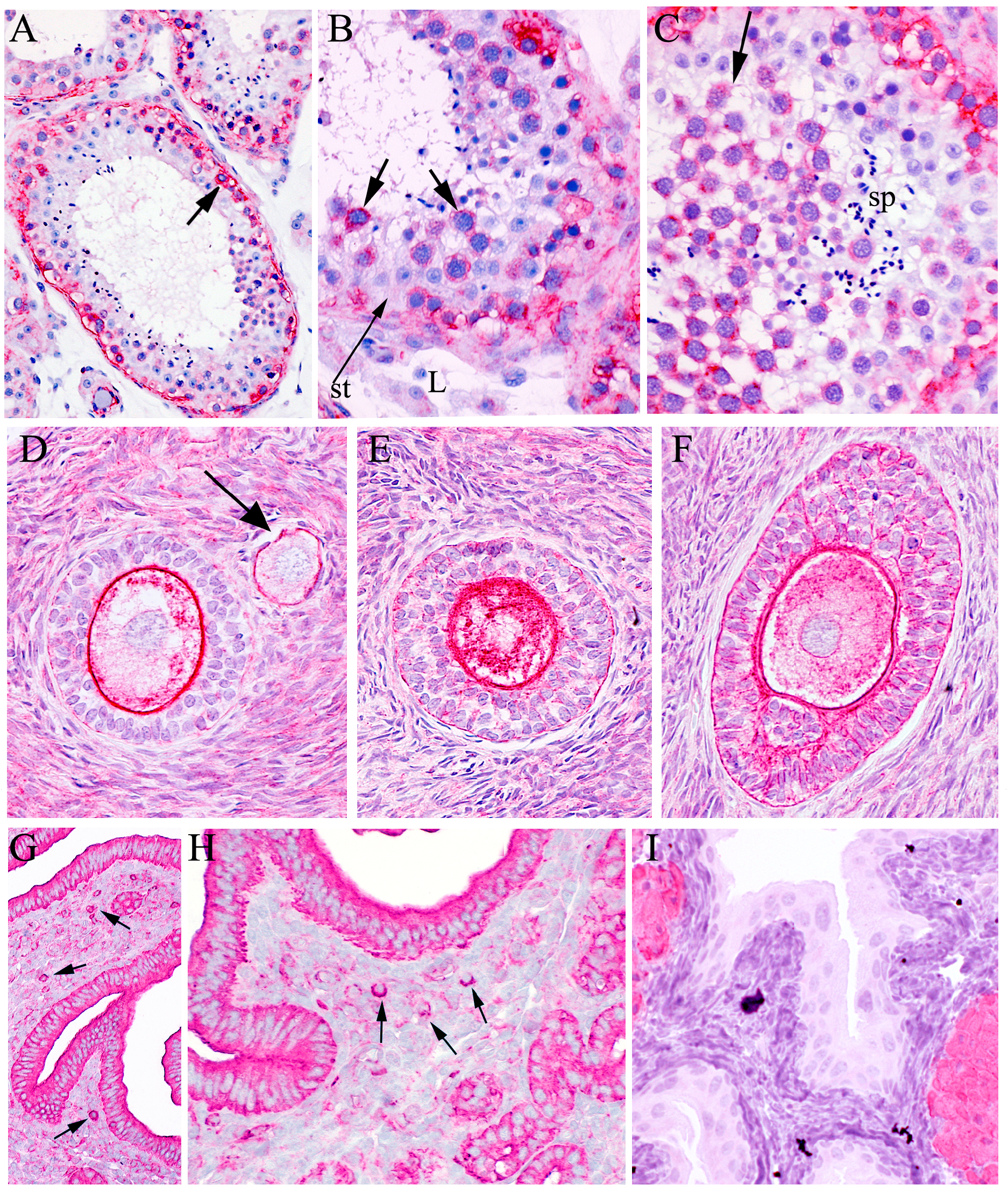
**A-C)** Human seminiferous tubules show strongly positive spx1 in diploid basal primary spermatogonia (as arrow in A). Secondary and meiotic spermatocytes are also positive as shown in B, but less intense (arrows) whereas spx1 is very low in almost mature sperm, Sertoli cells (st arrow), and interstitial Leydig cells (L) are only faintly positive. In C, mature sperm (sp) with negligible cytoplasm. show no spx1. **D-F)** human ovarian cells show increasing spx1 in the cytoplasm of maturing oocytes. In A, a less mature oocyte (arrow) in a unilayered follicle has a thin positive zona pellucida and weak cytoplasmic spx1 signal. Note the stronger cytoplasmic and zona pellucida signal in the adjacent multilayered follicle. As the follicle develops the spx1 signal becomes very intense and the inner plasma membrane of the egg is very intense in (E) and the more developed follicle (F). F shows layered ganulosa cells now have a stronger spx1 signal. **G&H**) Vagina of mouse shows intense epithelial and glandular spx1 in parallel sections with no pretreatment (G) and with citrate-heat unmasking (H). Most submucosal cells show little if any signal except for a few intensely stained round cells at arrows that are probably macrophages. **I)** In contrast, epithelium of mouse fallopian tube shows little spx1 while smooth muscle internal control is strongly positive.

Developing human oocytes showed an increasing expression of spx1 with maturation (Fig. 4D to F). In D, a small early unilayered follicle and an adjacent multilayered follicle are both highlighted by a strongly positive zonal pellucidum. The smaller ovum is cut through its center that includes its nucleus, and it displays only a few small positive cytoplasmic puncta over a diffuse weak background signal. In contrast, the larger mono-to-bilayered follicle shows strong punctate spx1 aggregates at the periphery, whereas outer follicular cells are negative or minimally positive. This intense peripheral staining pattern in the oocyte is also obvious in section 4E through a pole of a larger egg that does not include the nucleus. As the secondary follicle further develops, the ovum contains a rim of spx1 around the nucleus as well as strong punctate signals with a preference for the peripheral cytoplasm, possibly favoring one pole of the cell (4F); membranes between follicle layers are also now clearly positive. In sum, it is apparent that ova are actively producing spx1 from their SPX1.8 circles. It is not known how these circular elements originally entered germline cells but one likely scenario, given the plethora of environmental Acinetobacter that inhabit the gut microbiome, would involve endosymbiosis and sequestration by myeloid cells in the gut with subsequent transport and exchange of viral segments to oocytes in the early mammalian evolution.

The various epithelial cells sampled above have demonstrated very distinct and specific patterns of spx1 in highly differentiated tissues and cells. There were four different patterns of positive spx1 concentration. In some cell types, even those limited to the GI tract, the cytoplasm was ***a***) strongly and diffusely positive (as in chief and parietal cells of the stomach), ***b***) polarized (as in basal colonic cells), or ***c***) very weak (as in superficial gastric epithelium). Crypt cells in the stomach also showed only ***d***) a punctate plasma membrane signal, a feature even more strongly apparent in squamous skin cells. Fig. 4G & H show a mouse uterus with one of the most intensely stained epithelia observed in studies here. The endometrial submucosa was negative except for a few cells, probably macrophages (arrows). In contrast to intensely positive uterine epithelium, the fallopian tube epithelium (Fig. 4I) shows no spx1 signal while the smooth muscle in this section shows its typically strong spx1 signal. Thus the morphological expression pattern of spx1 protein is exquisitely tailored to distinguish subtle varieties of epithelial differentiation. Such features are likely to be useful for diagnostic and prognostic evaluation of epithelial neoplasias.

A summary of spx1 expression in various tissues and cell types of mice, rats and humans is presented in Table 1. Several tissues from Syrian Golden Hamsters and guinea pigs showed the same basic patterns of spx1 expression.

**Table I:**
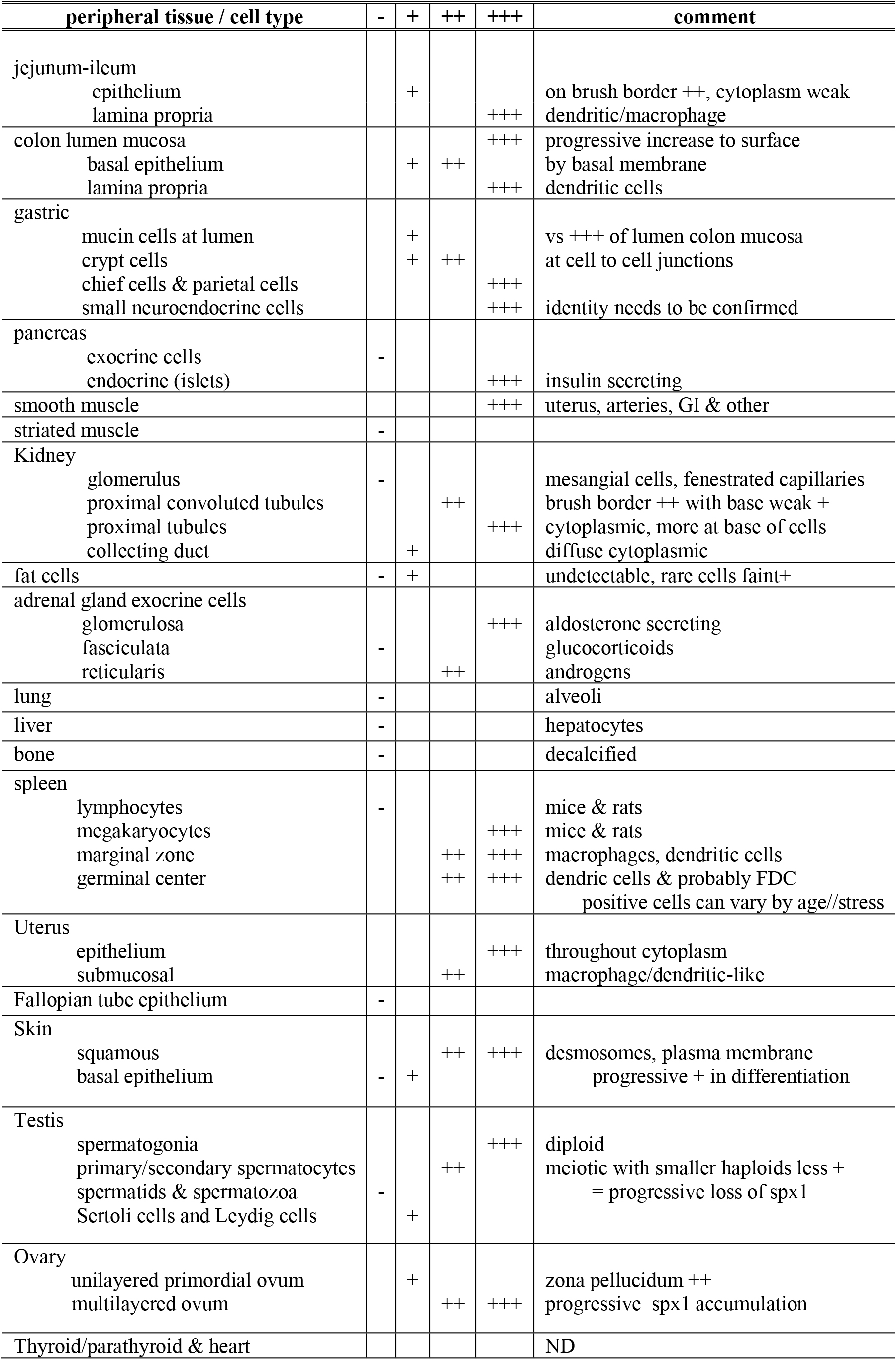
summary of spx1 signals in different tissues and cell types.

## DISCUSSION

SPHINX circular DNAs were discovered as a serendipitous offshoot from work on TSE infectious particles (21) and they open a glimpse into an unanticipated world of antediluvian viruses that can be incorporated and persist in mammalian cells. SPX1.8 DNA exists in a protected form in the cytoplasm and is inherited maternally, as evidenced here by its very strong expression in oocytes but not mature sperm. What are the function(s) of spx1 that would assure the conservation of its parental circular DNA in evolution? Some of its critical functions are suggested by its expression pattern in different cell types and tissues. The complete obliteration of signal by the antigenic peptide, and the identification of the same 41kd protein by two independently produced antibodies to REP, strongly support the SPHINX DNA origin of the spx1 signal here.

Remarkably, the spx1 protein accumulates only in certain types of cells, and these preferences were not anticipated by lineage. Intense spx1 in mesenchymal smooth muscle cells that contract as pulsatile syncytia, as in arteries and the GI tract, suggest spx1 functions in coordinated synchronous pulsation, a function that may share features with interactive neuronal synapse communications. Unlike smooth muscle, spx1 was not detected in striated muscle where fused multinucleated cells function together as discrete bundles. Indeed, unrestricted bundle to bundle excitation in striated muscle would lead to imprecise and uncoordinated voluntary neural control. A role for spx1 in cell to cell communication and strong junctional membrane connections in some epithelial cells, such as intestinal crypt cells, and the adherent desmosomes of squamous skin cells that differentiate into protective sheets, further emphasizes communicative functions.

A common stem cell lineage did not predict the strong spx1 expression in hematopoietic megakaryocytes, or its absence in follicular T cells. Epithelial cells of the pancreas also showed major spx1 differences, with negative exocrine cells adjacent to strongly positive insulin producing islet cells (31). This may be the result of spx1 repression, and methylation of SPX1.8 is one mechanism for repression that may be utilized. SPX1.8 expression may also require the presence of another circular DNA in the cell. SPX1.8 transcriptional activity and RNA synthesis has been complemented and enhanced by coinfection with a second SPHINX isolate from cows. On the other hand, the absence of spx1 signal in mature sperm signifies the elimination of SPX1.8 circles in specific negative cell types. Cells that lacked spx1 signals included major mammalian cell types such as lung alveoli, hepatocytes and bone. Although the loss of cytoplasmic mtDNA in sperm is well established and mtDNA can vary in different cell types, we are unaware of somatic cells normally lacking mtDNA. Attempts to evaluate SPX1.8 using Phi29 polymerase for in-situ amplification did not produce a meaningful signal in paraffin embedded formalin fixed archival tissue sections. These highly cross-linked preparations are unlikely to allow penetration or processivity of the enzyme and they probably also contain highly fragmented DNA that would further limit rolling circle amplification. Other approaches, such as PCR of cytoplasmic DNA from purified tissue-specific cell types, may help to resolve the copy number of SPX1.8 in fully mature cell types. However, this is not a trivial task given the positive vascular and smooth muscle cells present in all tissues. On the other hand, hematopoietic cells such as T cells and macrophages can be cleanly sorted for quantitative PCR.

Other tissues analyzed here extended quantitative difference in spx1 expression to extremely different patterns of spx1 localization in closely related cell types. Kidney proximal tubule cells with only subtle differences in absorption functions were apparent: proximal convoluted tubules were strongly stained only on the brush border and at the base. In contrast, the other proximal tubules were intensely positive throughout their cytoplasm. Even more convincing for the exquisite selection of spx1 expression were the marginally different layers of the adrenal cortex. Adrenal glomerulosa cells that secrete aldosterone for Na+ retention showed strong spx1 signals throughout their cytoplasm. In contrast, glucocorticoid secreting fasciculata cells were negative. Yet these cells are very closely related and cortical glomerulosa cells can reversibly change into fasciculate cells both morphologically and functionally simply by the application hormones such as ACTH (33). This reinforces transcriptional and/or translational controls, rather than loss of SPX1.8 that may be restricted to sperm. Exquisite differences in the pattern of spx1 expression were also found in epithelial cells of the GI tract. Secretory cells, such as gastric chief and parietal cells were strongly positive, but not adjacent mucous neck and surface cells. A very different pattern of spx1 labelling was also demonstrated in adjacent crypt cells of the stomach where only cell-to-cell plasma membrane signals were prominent. Together these studies underscore an acquired evolutionary symbiotic synchrony with distinctive function(s) for spx1 that are preserved in specific cell types of mammalian tissues.

Other ongoing spx1 studies here have revealed both variable spx1 expression in melanomas and glioblastomas, and both can contain positive and negative cells. Spx1 expression thus might be of value in staging or predicting the behavior of particular subsets of malignant human neoplastic cells. Clearly there is a lot more to be learned about SPHINX DNAs and related prokaryotic circular sequences, especially in terms of functional contributions that favor their preservation and expression. Neither of these realms can be adequately addressed by the present in-situ studies. Nevertheless, the current observations introduce a new world of ancient endosymbiotic prokaryotic viral elements that exist in mammals in an episomal state, and that can be maintained by maternal inheritance during evolution. More detailed evaluations of the functions of spx1, especially with reference to human aging, neurodegeneration, immunity, and neoplastic transformation can offer new insights into the covert world viruses and their unappreciated symbiotic exchanges that connect us to a changing environment. With respect to AD and other late onset neurodegenerations, it will also be of interest to find if SPHINX DNAs act as collaborators for the persistence and/or expression of common latent infections.

## Acknowledgements

I thank Reynold Spector for his sage suggestions on the manuscript, Brian West for his help with cell evaluations on the GI tract, Alexander Vortmeyer and Harvey Kliman for human testis and ovarian tissue sections, and Laertes Manuelidis for providing human skin sections.

